# Generation of a recombinant GST-Tau antigen for indirect ELISA in evaluating Alzheimer’s disease vaccines

**DOI:** 10.1101/2025.09.18.676867

**Authors:** Diana I. Zamora-Loyarte, Karen León-Arcia, Gabriela Pérez-Leal, Heidi Quintero-Alvarez, Mailen López-Armenteros, Alexandra Sánchez-Quesada, Nathalí Nordelo-Ávila, Yaimeé Vázquez-Mojena

## Abstract

**Background:** Alzheimer’s disease is a progressive neurodegenerative disorder. Immunotherapy targeting Aβ and Tau proteins is a promising strategy, and accurate detection of anti-Aβ or anti-Tau antibodies is key to evaluating vaccine efficacy. Indirect ELISA is a sensitive method requiring high-quality coating antigens.

**Methodology:** A recombinant Tau antigen fused to glutathione S-transferase (GST) was designed as an ELISA coating antigen. The Tau gene was cloned into the pGEX-6p-1 vector and expressed in *Escherichia coli* BL21 cells. Protein expression was induced with 1 mmol/L isopropyl-β-D-thiogalactoside (IPTG), and the GST-Tau fusion protein was identified by Western blot using anti-Tau and anti-GST antibodies. The protein was solubilized with 3 mol/L urea and purified via glutathione affinity chromatography. Purity was confirmed by densitometry and the optimal coating concentration for ELISA was determined to be 2 μg/mL.

**Results:** A protocol was established to produce GST-Tau with adequate purity and yield for ELISA applications. The recombinant antigen retained immunogenic epitopes, recognized by anti-Tau antibodies. The standardized ELISA enabled sensitive detection of anti-Tau antibodies.

**Conclusions:** A robust method was developed to produce and optimize GST-Tau as a coating antigen for indirect ELISA. This approach facilitates reliable quantification of anti-Tau antibodies to evaluate Alzheimer’s vaccine candidates.

## Introduction

Alzheimer’s disease (AD) is a progressive neurodegenerative disorder whose prevalence increases dramatically with age, currently affecting approximately 51% of people aged 65 and older. ^(1)^ Projections indicate a significant rise in AD cases by 2050. ^(2)^ In Cuba, dementia affects 10.2% of those aged 65 and above, with an annual incidence of 21 per 1,000 in this age group, and the number of individuals living with dementia is expected to rise from 160,000 to 273,000 by 2040.^(3)^

Vaccination represents a promising approach for both the prevention and treatment of AD. By safely stimulating the immune system, vaccines can induce the production of antibodies that specifically target disease-related proteins such as Aβ or Tau. ^(4)^ Accurate measurement of antibody titers is crucial for evaluating the effectiveness of experimental vaccine candidates. Analytical systems that detect specific circulating antibodies in serum provide reliable confirmation of the immune responses following vaccination.

Enzyme-linked immunosorbent assay (ELISA) is widely used to evaluate humoral responses in animal models and clinical practice. ^(5)^ While commercial indirect ELISA (iELISA) kits are available, developing and standardizing an in-house iELISA protocol can be more economical and flexible. Standardization involves optimizing key parameters such as the concentration of the coating antigen, detection antibody, and sample dilution. ^(6, 7)^ High purity and stability of the coating antigen are essential for effective immobilization on ELISA plates.

The glutathione-S-transferase (GST) fusion-protein system is commonly employed in vaccine research because GST-tagged proteins are more soluble and can be efficiently purified via glutathione affinity chromatography. ^(8, 9)^ The GST tag also facilitates the immobilization of the antigen on ELISA plates, exposing relevant epitopes for antibody detection. ^(10)^ In this study, we produced a GST-Tau fusion protein as an alternative coating antigen ^(12)^ to evaluate humoral responses generated by Alzheimer’s vaccine candidates.

## Materials and Methods

### Induction and expression of the GST-Tau protein

A synthetic gene encoding tau epitopes with the sequence:*atggccgaaccgcgccaagaatttgaggtgatggaggaccatgccggtacctatggtttaggtgatcgtaaggatcaaggtggctat accatgcatcaagatcaagaaggtgataccgatgctggtctgaaagaaagtccgctgggtggcggcggcagctttaacaactttaccgtgagcttttggctgcgtgtgccgaaagtgagcgccagccatctggaaggtggcggtggcagcagtaaatgcggctctttaggcaatattcaccataaaccgggcggtggtcaagttgaagtgaagagcgagaaactggacttcaaagaccgcgtgcagagcaaaatcggctctttagacaacattacccatgtgccgggtggcggcaacaagaaaattgagacccataaactgaccttccgcgaaaacgccaaagccaaaaccgatcatggcgccgagatcgtgtataaagaaccggtggtgagcggcgatacagaaccgcgtcatctgagcaatgttagcagcaccggcagcatcgatatggtggatgagcctcagctggccactttagccgacgaagttagcgcctctttagccaaacaaggtctgggatcccgggca* (13) was cloned into the pGEX-6p-1 vector to generate the GST-Tau fused protein. *E. coli* BL21 (DE3) cells were transformed with the recombinant plasmid and grown overnight at 37°C in Luria Bertani (LB) medium with ampicillin (100 μg/mL). The culture was expanded in 500 mL fresh LB-ampicillin (100 μg/mL) and incubated at 37°C to an optical density 600 nm (OD600) of 0.6. Protein expression was induced with 1 mmol/L IPTG, followed by incubation at 28 °C for an additional 3 hours.

### GST-Tau protein identity verification by Western blot analysis

Samples from the induced culture were mixed with 2X loading buffer, heated at 95 °C for 5 min, and separated by 12% SDS-PAGE. Proteins were transferred to a nitrocellulose membranes and blocked with 5% milk in PBS-0.05 mol/L Tween20 (PBS-T) for 1 hour at 37°C. Membranes were incubated for 1 hour at 37 °C with either an anti-goat GST (CIGBSS, Cuba) (1:3,000) or an anti-mouse Tau monoclonal antibodies (EMD Millipore, MAB2239) (1:5,000) diluted in 2% skimmed milk. After washing with PBS-T, membranes were incubated for 1 hour at 37 °C with the appropriate HRP-conjugated secondary antibody (goat-anti-mouse for Tau detection (SIGMA, A3438) or rabbit-anti-goat for GST detection (CIGBSS, Cuba), each diluted 1:5,000. Bands were visualized using diaminobenzidine as substrate for five minutes. The reaction was terminated decanting the substrate and rinsing the membrane with distilled water.

### Solubilization of GST-Tau protein

Cells were harvested by centrifugation at 5,000 g, 4°C for 15 min, and resuspended in 4 mL of lysis buffer (50 mmol/L Tris pH 8.0, 1 mmol/L PMSF, and 5 mmol/L DTT) per 100 mL culture. Lysis was performed with 1 mg/mL lysozyme and incubating on ice for 30 min, followed by centrifugation at 10,000 g, 4°C for 10 min. The insoluble fraction was resuspended in 8 mL of solubilization buffer (3 mol/L de Urea, 25 mmol/L de triethanolamine, 1 mmol/L de EDTA pH 8.0, 5 mmol/L de DTT, 1 mmol/L PMSF, and 150 mmol/L NaCl) and incubated 10 min at 4°C. Soluble proteins were recovered by centrifugation at 10,000g, 4°C for 10 min.

### Purification of GST-Tau protein

GST-Tau protein was purified by affinity chromatography using immobilized glutathione resin. The soluble fraction was incubated with the resin for 2 hours at 4°C, washed with buffer (50 mmol/L Tris, pH 8.0, 1 mmol/L PMSF, and 5 mmol/L DTT), and eluted with 10 mmol/L reduced glutathione in buffer containing 1 mmol/L PMSF and 150 mmol/L NaCl for 10 minutes. Five elution fractions were collected, monitored by absorbance at 280 nm, and stored at −70°C.

### Standardization of the coating antigen concentration

ELISA plates were coated overnight at 4°C with 100 μL/well of GST-Tau protein at concentrations from 1.25 to 5 µg/mL. After coating, the plates were washed four times with 200 μL/well of 0.05% PBS-Tween 20 and then blocked with 200 µl 5% skimmed milk in PBS for 1 hour at room temperature to prevent non-specific binding.

Anti-Tau antibody was diluted in PBS containing 2% skimmed milk at concentrations of 1:500, 1:1,000; 1:2,000; 1:4,000; 1:8,000; 1:16,000; and 1:32,000. A volume of 100 µl per well was added and incubated at 37 °C for 1 hour. After washing, HRP-conjugated mouse anti-IgG secondary antibody (SIGMA, A9917) (1:5,000) was added and incubated at 37 °C for 1 hour. O-Phenylenediamine dihydrochloride peroxidase (OPD) substrate was added to each well, and color development was allowed to proceed for 30 minutes before the reaction was stopped with 2N H_2_SO_4_. Absorbance at 492 nm was measured and plotted against the anti-Tau antibody concentration, and analyzed using a 4PL sigmoidal fit. Non-linear regression (R^2^) values of these curves were compared, and the GST-Tau coating concentration that produced the highest R^2^was selected as optimal for the ELISA plates.

### Detection of anti-Tau antibodies in immunized mice

To demonstrate the functionality of GST-Tau as a coating antigen, an experimental assay was performed in male C57BL/6 wild-type mice (n=6/group). All the experiments were performed in accordance with legal and institutional guidelines and were carried out under ethics, consent, and permissions of the Research Ethical Committee of Cuban Center for Neuroscience.

The animals were divided into three experimental groups administered 25, 50 or 100 μg/mL doses of the PK-Tau vaccine candidate (13), while a control group received 100 μL of Tris 50 mmol/L. All antigen formulations were adsorbed onto aluminum hydroxide as an adjuvant. Immunization was delivered subcutaneously every two weeks for a total of five doses. Serum samples were collected pre-immunization at each timepoint for antibody analysis.

Anti-Tau antibody levels were assessed via iELISA using plates coated with 2 μg/mL GST-Tau fusion protein, following the established protocol. Quantification measurements were performed in duplicate with appropriate controls.

Statistical analysis was conducted using GraphPad Prism 9.5.1 (GraphPad Software, Inc.) with statistical significance defined as p < 0.05. Data are presented as mean ± standard error of the mean. Comparisons of anti-Tau antibody concentrations among the experimental groups were performed using multivariate analysis of variance (MANOVA).

## Results and Discussion

### Expression and immunodetection of GST-Tau protein

After IPTG induction, *E. coli* BL21 (DE3) cells expressing pGEX6p-1-Tau displayed a distinct protein band at approximately 48 kDa, which was absent in untransformed controls (Fig. 1a). This aligns with the expected molecular weight of the GST-Tau antigen, as previously estimated by *in silico* analysis and confirmed by densitometry, showing comparable expression levels to those reported by Pérez. ^(12)^ Western blot analysis revealed that both anti-GST and anti-Tau antibodies recognized this band (Fig.s 1b and c, respectively), confirming the identity of the fusion protein.

**Fig. 1.**
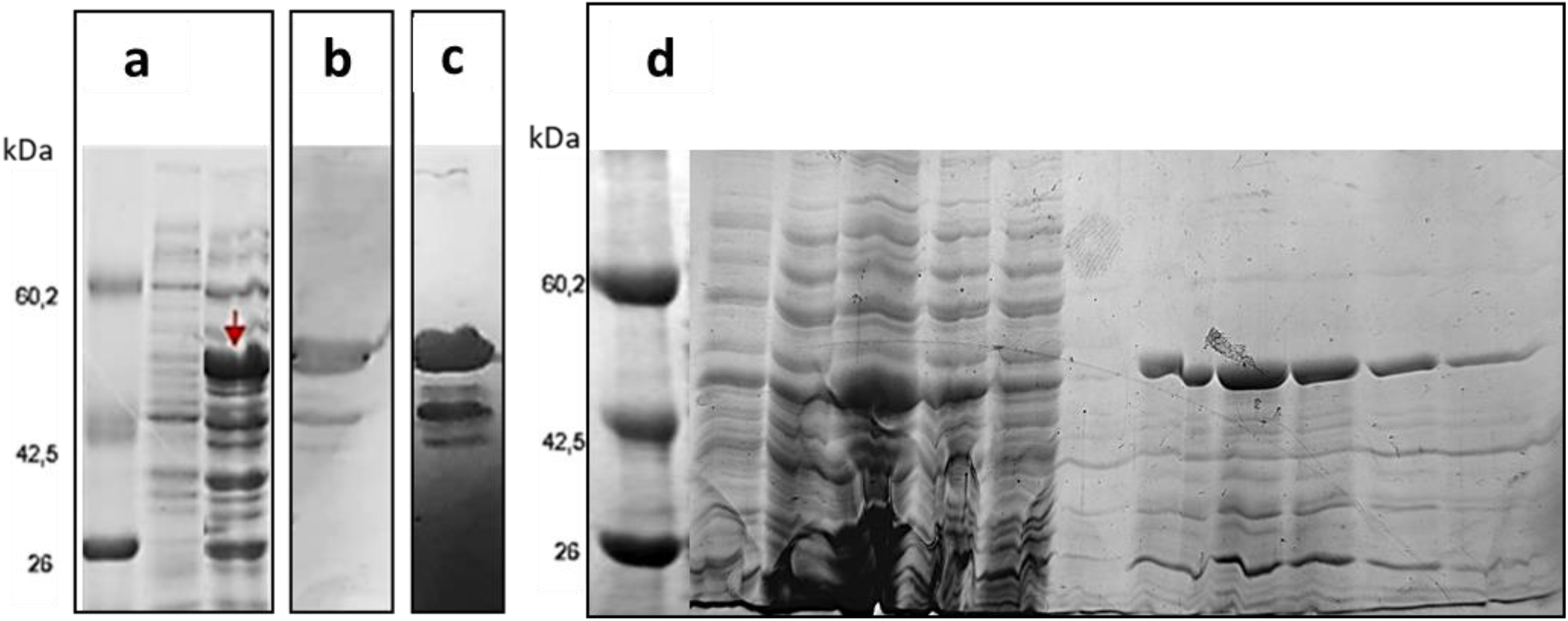
Expression, immunodetection, and purification of recombinant GST-Tau fusion protein from E. coli BL21 (DE3): **(a)** SDS-PAGE analysis of total protein extracts from *E. coli* BL21 (DE3) cells transformed with the pGEX6p-1-Tau plasmid. Lane 1: molecular weight marker; Lane 2: uninduced control; Lane 3: IPTG-induced sample showing a distinct band at ~48 kDa corresponding to the GST-Tau fusion protein (red arrow). **(b)** Western blot of the induced sample probed with anti-GST antibody, confirming the presence of the GST tag in the expressed fusion protein. **(c)** Western blot probed with anti-Tau antibody, confirming the Tau component of the fusion protein. Lower molecular weight bands indicate possible degradation products, while higher molecular weight bands suggest multimer formation. **(d)** SDS-PAGE analysis of fractions collected during purification of GST-Tau by glutathione affinity chromatography. Lane 1: molecular weight marker; Lane 2 and 3: supernatant and pellet after cell lysis with lysozyme; Lane 4 and 5: supernatant and pellet after 3 mol/L urea solubilization, respectively; Lane 6: supernatant after cell lysis and binding to resin; Lane 7column wash; Lanes 8-12: sequential elution fractions containing purified GST-Tau protein. The predominant band at ~48 kDa corresponds to the GST-Tau fusion protein; additional bands indicate possible aggregates or degradation products

However, additional bands of lower molecular weight were observed, indicating partial proteolytic degradation of the chimeric antigen during expression (Fig. 1b and c). This is likely due to the use of high IPTG concentrations, which can stress *E. coli* and promote proteolysis. ^(14)^ Future optimizations could include reducing IPTG concentration, lowering induction temperature, or using protease-deficient strains to improve protein integrity. ^(12, 15, 16)^

High molecular weight bands were also evident (Fig. 1c), indicating the formation of Tau multimers, as the anti-Tau antibody detected proteins with higher molecular weights than expected (Fig. 1c). This is consistent with previous reports that Tau proteins, particularly those with intact N-terminal regions can form dimers and trimers under denaturing conditions. ^(17, 18)^ Adjusting the concentration of reducing agents like β-mercaptoethanol or ditiothreytol (DTT), or modifying sample heating times during SDS-PAGE may help minimize aggregation. ^(19)^ Additionally, lower induction temperatures could reduce aggregation by minimizing hydrophobic interactions. ^(14)^

### Purification of the GST-Tau protein

GST-Tau was efficiently solubilized using lysozyme in combination with 3 mol/L urea (Fig. 1d, lanes 3 and 4). However, some antigen loss was detected in the flow-through fractions (Fig. 1d, lane 5), likely due to partial denaturation reducing GST’s affinity for the glutathione resin. Although urea concentrations of up to 4 mol/L are generally compatible with GST binding, ^(20)^ further optimization of solubilization and binding conditions may enhance the overall yield. Schäfer *et al.* have demonstrated that urea concentrations as high as 4 mol/L do not impair GST binding to immobilized glutathione during cell lysis. Additionally, the inclusion of 150 mmol/L NaCl was based on previous studies that optimized solubilization of inclusion bodies. ^(10)^

Elution fractions showed a predominant ~48 kDa band (Fig. 1d, lanes 7-11), with a purity of ~85% as estimated by densitometry. Minor bands likely represent degradation products or aggregates. The final concentration of GST-Tau was 0.11 mg/mL, sufficient for ELISA plate coating.

### Standardization of coating antigen concentration for iELISA

Although complete purity is not essential for ELISA coating antigens, ^(21, 22)^ optimizing antigen concentration is critical. Low concentrations may yield weak signals, while excessive antigen can cause monolayer formation and reduced sensitivity. ^(23)^

Regression analysis of the ELISA data (Fig. 2) showed that 2 μg/mL GST-Tau provided the highest R^2^value (0.9966), indicating optimal sensitivity. Lower (1.25 μg/mL, R^2^= 0.9940) and higher (5 μg/mL, R^2^= 0.9933) concentrations were less effective, likely due to under- or over-coating effects. These results are consistent with literature recommendations for antigen concentrations between 1 and 10 μg/mL. ^(22, 24)^

**Fig. 2.**
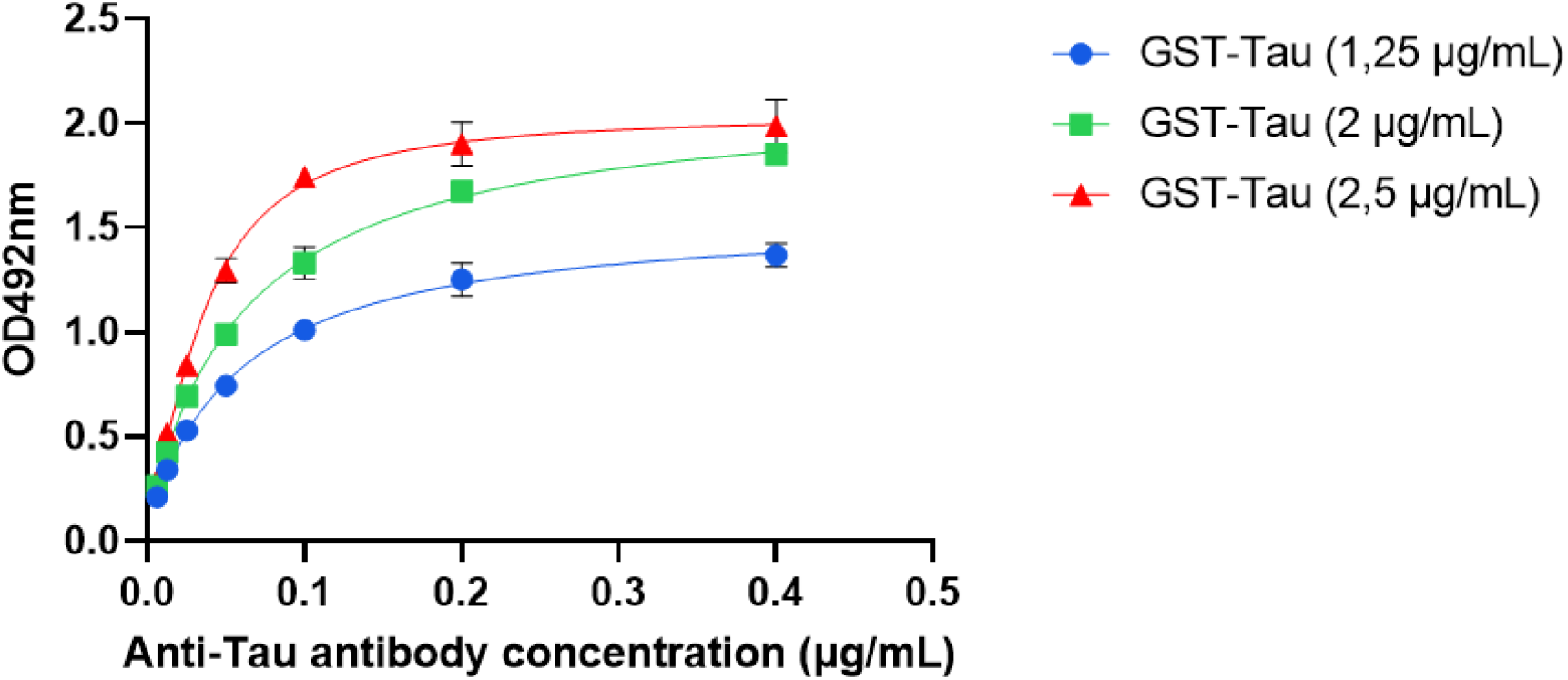
Optimization of GST-Tau antigen concentration for ELISA plate coating based on 4PL sigmoidal fit of antibody detection. Non-linear regression analysis of optical density (OD492nm) versus anti-Tau antibody concentration for ELISA plates coated with three different concentrations of GST-Tau antigen (1.25, 2, and 2.5 μg/mL). Colored lines represent the non-linear fit for each antigen concentration: blue (1.25 μg/mL), green (2 μg/mL), and red (2.5 μg/mL). The highest R^2^value observed at 2 μg/mL indicates this is the optimal antigen concentration for sensitive antibody detection in indirect ELISA

### Detection of anti-Tau antibodies in immunized mice

The suitability of the in-house iELISA, using 2 μg/mL GST-Tau as the coating antigen, was validated by quantifying the antibody response in mice immunized with an anti-Tau vaccine candidate. The assay detected significantly elevated levels of specific anti-Tau antibodies in the sera of vaccinated mice compared to placebo controls, demonstrating a clear dose-dependent response, with the highest antibody titers observed in the group receiving 100 μg of the vaccine (Fig. 3).

**Fig. 3.**
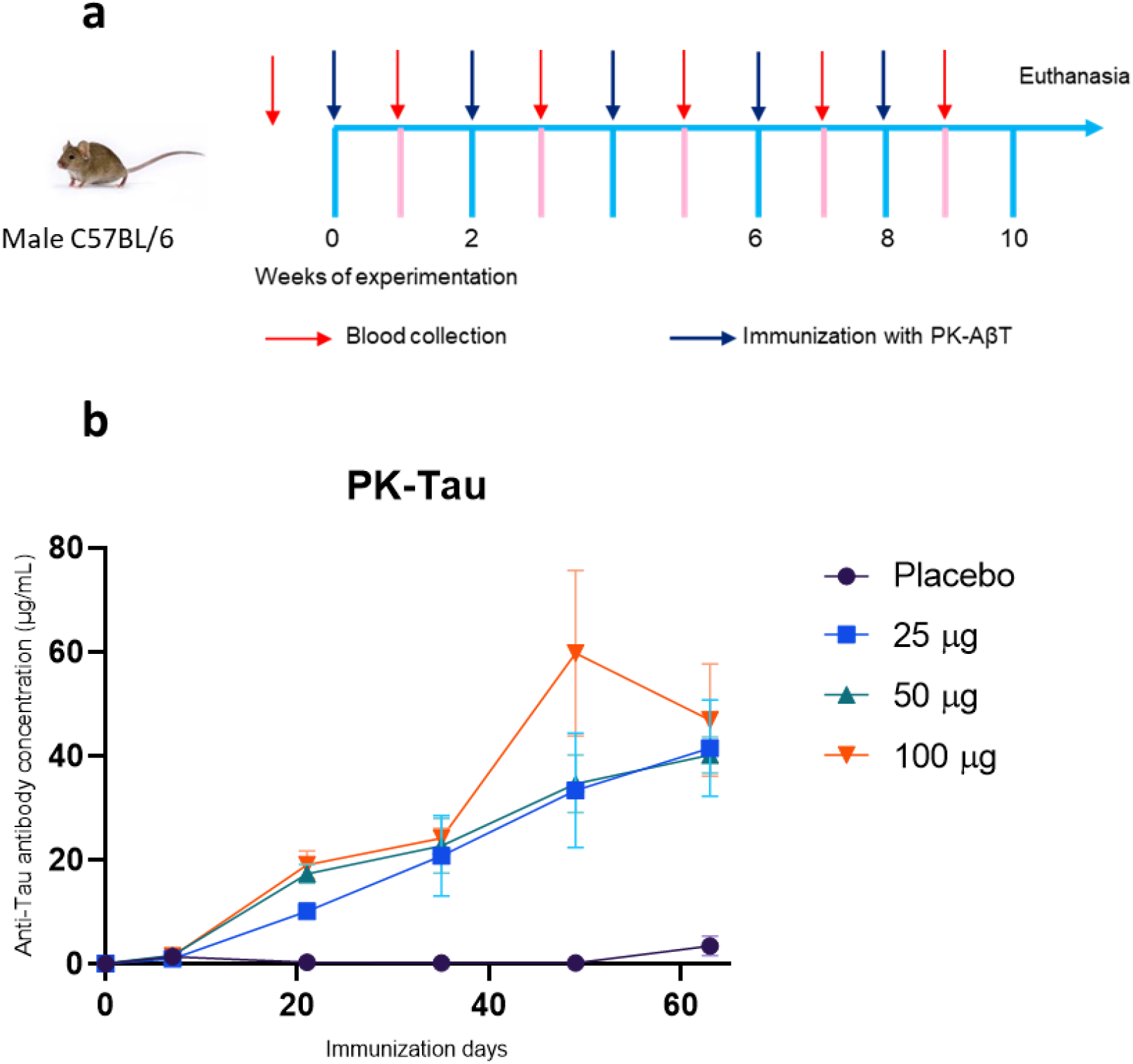
Immune response in C57BL/6 mice vaccinated with an anti-Tau formulation. **(a)** Schematic outlining the immunization (Created with BioRender.com). **(b)** Concentration of anti-Tau specific antibodies. Data are presented as mean ± SEM

These results demonstrate that a GST-Tau fusion protein can be efficiently produced and effectively used as a coating antigen for indirect ELISA in Alzheimer’s vaccine research. This method is both cost-effective and adaptable, making it especially suitable for laboratories with limited resources.

Nonetheless, further optimization could enhance protein yield and integrity. Strategies such as lowering IPTG concentrations, employing auto-induction media, or reducing expression temperatures may help minimize proteolytic degradation and aggregation. Additionally, incorporating alternative solubilization methods, such as the use of mild detergents or molecular chaperones, could further reduce aggregation and improve antigen quality.

Despite these areas for improvement, the standardized GST-Tau antigen developed in this study is now ready for use in evaluating antibody responses in animal models. Moreover, the methodology described here can be readily adapted for the production and standardization of other antigens relevant to neurodegenerative disease research.

## Conclusions

We have successfully produced a GST-Tau fusion protein suitable for use as a coating antigen in indirect ELISA assays. The antigen was recognized by specific anti-Tau antibodies, confirming the preservation of relevant immunogenic epitopes. An optimal coating concentration of 2 μg/mL was established, providing a robust platform for evaluating anti-Tau antibody responses in preclinical studies of Alzheimer’s vaccines. Further optimization of expression and purification protocols may enhance yield and antigen quality. Validating the ELISA by determining the coefficients of variation of robustness and specificity assays will improve the anti-tau antibody detection system. This will broaden its applicability in neurodegenerative disease research.

## Statements and Declarations

## Acknowledgments

Karen León-Arcia would like to thank the Mexican Secretariat of Science, Humanities, Technology and Innovation (Secihti) for grant No. 2007712.

## Declaration of Generative AI and AI-assisted technologies in the writing process

During the preparation of this work the author(s) used Grammarly in order to improve readability and language, and check grammar and spelling. After using this tool/service, the author(s) reviewed and edited the content as needed and take(s) full responsibility for the content of the publication.

## Author Contributions

Conceptualization: Karen León-Arcia, Heidi Quintero-Alvarez Data curation: Diana I. Zamora-Loyarte, Yaimeé Vázquez-Mojena

Formal Analysis: Diana I. Zamora-Loyarte, Karen León-Arcia, Gabriela Pérez-Leal

Funding acquisition: Karen León-Arcia

Investigation: Diana I. Zamora-Loyarte, Karen León-Arcia, Gabriela Pérez-Leal, Heidi Quintero-Alvarez, Mailen López-Armenteros, Alexandra Sánchez-Quesada, Nathalí Nordelo-Ávila

Metodology: Karen León-Arcia

Project administration: Karen León-Arcia

Resources: Karen León-Arcia

Software:

Supervission: Karen León-Arcia

Validation: Karen León-Arcia, Heidi Quintero-Alvarez, Yaimeé Vázquez-Mojena Visualization:

Writing – original draft: Diana I. Zamora-Loyarte

Writing – review & editing: Karen León-Arcia, Gabriela Pérez-Leal, Heidi Quintero-Alvarez, Mailen López-Armenteros, Alexandra Sánchez-Quesada, Nathalí Nordelo-Ávila, Yaimeé Vázquez-Mojena

## Ethics approval

All the experiments were performed in accordance with the Declaration of Helsinki and institutional guidelines. Approval was granted by the Research Ethical Committee of Cuban Neuroscience Centre.

## Consent to publish

All authors attest they meet the ICMJE criteria for authorship and approve the submitted manuscript.

## Availability of data and materials

The analyzed data sets generated during the current study are available from the corresponding author on reasonable request.

## References

1. Aguzzoli CS, et al. World Alzheimer Report 2024 Global changes in attitudes to dementia Contributors: Survey translators [Internet]. 2024. Available from: https://www.alzint.org/resource/world-alzheimer-report-2024/ Accessed 2025 Mar 18.

2. Cummings J, et al. Alzheimer’s disease drug development pipeline: 2024. Alzheimer’s Dementia Transl Res Clin Interv. 2024;10(2):e12465. doi:10.1002/trc2.12465.

3. Llibre-Rodríguez J, Gutiérrez-Herrera R. Avances y desafíos en la investigación de la enfermedad de Alzheimer. Anales de la Academia de Ciencias de Cuba [Internet]. 2024;14(1). Available from: https://revistaccuba.sld.cu/index.php/revacc/article/view/1483 Accessed 2025 Apr 12.

4. Vukicevic M, et al. An amyloid beta vaccine that safely drives immunity to a key pathological species in Alzheimer’s disease: pyroglutamate amyloid beta. Brain Commun. 2022;4(1). doi:10.1093/braincomms/fcac022.

5. Liu N, Liang X, Chen Y, Xie L. Recent trends in treatment strategies for Alzheimer’s disease and the challenges: A topical advancement. Ageing Res Rev. 2024;94:102199. doi:10.1016/j.arr.2024.102199.

6. Kohl TO, Ascoli CA. Indirect immunometric ELISA. Cold Spring Harb Protoc. 2017;2017(5):396–401. doi:10.1101/pdb.prot093708.

7. Minic R, Zivkovic I. Optimization, Validation and Standardization of ELISA. In: Norovirus. IntechOpen; 2020. doi:10.5772/intechopen.94338.

8. Asaduzzaman M, et al. The Technique for Excessive-Results of Recombinant Protein Purification led to GST-tag Affinity Chromatography: A Review. J Microb Biochem Technol. 2023;15:584. doi:10.35248/1948-5948.23.15.584.

9. Harper S, Speicher DW. Purification of Proteins Fused to Glutathione S-Transferase. In: Methods in Molecular Biology. Vol 681. Humana Press Inc.,, 2011. p. 259–280. doi:10.1007/978-1-60761-913-0_14.

10. Shendge AA, D’Souza JS. Strategic optimization of conditions for the solubilization of GST-tagged amphipathic helix-containing ciliary proteins overexpressed as inclusion bodies in E. coli. Microb Cell Fact. 2022;21(1):258. doi:10.1186/s12934-022-01979-y.

11. Zhu J, Song H, Le G. A review of chimeric proteins/enzymes. BIO Web Conf. 2024;111:1017. doi:10.1051/bioconf/202411101017.

12. Perez G. Optimización de la expresión en Escherichia coli de tres antígenos quiméricos fusionados a Glutatión-S-Transferasa, propuestos como candidatos vacunales para la enfermedad de Alzheimer [tesis de licenciatura]. Universidad de La Habana; 2021.

13. Quintero H. Diseño y obtención de antígenos quiméricos multiepitópicos basados en regiones inmunogénicas de las proteínas Aβ y tau como candidatos a antígenos vacunales contra Enfermedad de Alzheimer [tesis de maestría]. Universidad de La Habana; 2021.

14. Lozano G, et al. Impact of the Expression System on Recombinant Protein Production in Escherichia coli BL21. Front Microbiol. 2021;12. doi:10.3389/fmicb.2021.682001.

15. González A, Fillat MF. Aspectos metodológicos de la expresión de proteínas recombinantes en Escherichia coli. Rev Educ Bioquim. 2018;37(1):14–27.

16. Pouresmaeil M, Azizi-Dargahlou S. Factors involved in heterologous expression of proteins in E. coli host. Arch Microbiol. 2023;205(5). doi:10.1007/s00203-023-03541-9.

17. Zhang X, Wang J, Zhang Z, Ye K. Tau in neurodegenerative diseases: molecular mechanisms, biomarkers, and therapeutic strategies. Transl Neurodegener. 2024;13(1):40. doi:10.1186/s40035-024-00429-6.

18. Magalhães AD, et al. Large-scale seroepidemiology uncovers nephrological pathologies in people with tau autoimmunity [tesis doctoral]. University of Zurich; 2021. doi:10.1101/2021.11.24.21266833.

19. Mahmood T, Yang PC. Western blot: Technique, theory, and troubleshooting. North Am J Med Sci. 2012;4(9):429–34. doi:10.4103/1947-2714.100998.

20. Schäfer F, et al. Chapter Nine - Purification of GST-Tagged Proteins. In: Lorsch JR, editor. Laboratory Methods in Enzymology: Protein Part D. Methods in Enzymology, vol. 559. Academic Press; 2014. p. 127–39. doi:10.1016/bs.mie.2014.11.005.

21. Crowther J. The ELISA Guidebook. Vol 516. 2009. doi:10.1007/978-1-60327-254-4.

22. Mandiarote A, et al. Estandarización de ensayos inmunoenzimáticos (ELISA) para la cuantificación de anticuerpos IgG inducidos por una vacuna de vesículas de membrana externa de los serogrupos A y W 135 de Neisseria meningitidis [Internet]. Vaccimonitor Accessed 2025 Jun 5. Available from: https://www.finlay.sld.cu/vaccimonitor.htm

23. Choquehuanca JL, et al. Estandarización y validación de pruebas de ELISA tipo indirecto para la determinación de los niveles de anticuerpos IgG e IgE anti-leishmania, como método complementario para el diagnóstico y seguimiento de la respuesta al tratamiento. Revista CON-CIENCIA. 2019;7:39–54. Accessed 2025 June 5. Available from: http://www.scielo.org.bo/scielo.php?script=sci_arttext&pid=S2310-02652019000200005&nrm=iso

24. Pino Y, et al. Validación de un ensayo ELISA para la determinación de anticuerpos anti LPS de Vibrio cholerae. Vaccimonitor. 2003;12:11–17. Accessed 2025 June 5. Available from: http://scielo.sld.cu/scielo.php?script=sci_arttext&pid=S1025-028X2003000100002&nrm=iso

